# DIFFERENTIAL PHOTOSYNTHETIC RESPONSES TO GLUFOSINATE AMMONIUM IN TWO GRASS WEEDS: *Lolium multiflorum* AND *Echinochloa crus-galli*

**DOI:** 10.64898/2026.08.31.748356

**Authors:** Ornella Sabatini, Juana Villalba, Sebastián Simondi, María Martha Sainz, Gastón Quero

**Affiliations:** Ing. Agr., Cibeles S.A., Montevideo, Uruguay; Profesor Agregado, Departamento Protección Vegetal, Estación Experimental Dr. Mario A. Cassinoni, Facultad de Agronomía, Paysandú, Uruguay; Prof. Titular, Área de Matemática, Facultad de Ciencias Exactas y Naturales, Universidad Nacional de Cuyo (FCEN-UNCuyo), Mendoza, Argentina; Prof. Adjunto, Departamento de Biología Vegetal, Facultad de Agronomía, Universidad de la República, Montevideo, Uruguay

**Author notes:** Ornella Sabatini Rodríguez, Juana Villalba, Sebastián Ricardo Simondi, María Martha Sainz.

**Keywords:** Herbicide resistance, Chlorophyll fluorescence, Leaf senescence, Quantum yield PSII

## Abstract

**Background:** Weed control is one of the main challenges in agriculture today, particularly due to the increasing occurrence of herbicide-resistant populations. Among the most problematic species are *Lolium multiflorum* (L.) and *Echinochloa crus-galli* (L.) Beauv., for which glyphosate-resistant populations have been reported. In this context, glufosinate ammonium has emerged as an alternative for their control; however, its efficacy may vary depending on species and photosynthetic metabolism.

**Objective:** The objective of this study was to evaluate the differential sensitivity of ryegrass (C3) and barnyardgrass (C4) to ammonium glufosinate by analyzing physiological responses associated with leaf senescence and photosystem II activity.

**Methods:** Visual injury, chlorophyll fluorescence, and ammonium accumulation were assessed.

**Results:** Results revealed a differential response between species. Barnyardgrass exhibited earlier symptom onset and a greater reduction in the quantum yield of photosystem II (ΦPSII), whereas ryegrass showed a slower senescence process. These differences indicate a higher sensitivity of barnyardgrass to glufosinate ammonium, possibly associated with its C4 photosynthetic metabolism.

**Conclusions:** It is concluded that the effectiveness of glufosinate ammonium depends on the type of photosynthetic metabolism and on the ability of each species to cope with herbicide-induced oxidative stress. This information contributes to optimizing glufosinate ammonium use and to the development of management strategies aimed at delaying the evolution of herbicide resistance.

**Highlights:**

- Glufosinate efficacy differs between C3 ryegrass and C4 barnyardgrass.
- Barnyardgrass shows earlier symptoms and stronger PSII inhibition than ryegrass.
- Species differ in senescence and stress tolerance under glufosinate exposure.

## 1. Introduction

Weed control is one of the major challenges in agriculture today, particularly due to the increasing occurrence of herbicide-resistant populations. Grass weeds such as *Lolium multiflorum* and *Echinochloa crus-galli* are widely distributed in agricultural systems and can cause significant yield losses. In Uruguay, these species are among the most problematic weeds due to their highly competitive ability and the evolution of resistance to herbicides (García et al., 2025).

*Lolium multiflorum* (ryegrass) is a C3 grass species widely distributed in winter cropping systems, affecting crops such as wheat, barley, and canola, as well as fallow periods prior to summer crops. In Uruguay, it is one of the most frequent winter weeds and has shown a high adaptive capacity under intensive agricultural systems (Ríos et al., 2005; García et al., 2025). Since the first report of glyphosate-resistant populations in 2016, multiple resistance mechanisms have been documented, including resistance to graminicides and ALS-inhibiting herbicides (Marques et al., 2022).

On the other hand, *Echinochloa crus-galli* (barnyardgrass) is a C4 annual species that represents a major weed problem in summer crops and rice systems (Belgeri and Caulin, 2008; Mailhos and San Román, 2008; García et al., 2025). In Uruguay, resistant biotypes to several herbicide modes of action, including propanil and imidazolinones, have been reported, as well as cases of multiple resistance (Metzler et al., 2018; Marchesi and Saldain, 2019). Both species are characterized by high seed production, strong adaptability, and a high capacity to evolve resistance, making their management particularly challenging.

In this context, glufosinate ammonium becomes an alternative for the control of these weeds in different winter and summer crops. Glufosinate ammonium is classified as a non-selective herbicide (Shaner 2014), inhibits the enzyme glutamine synthetase, leading to ammonium accumulation, altered nitrogen metabolism, and foliar necrosis (Coetzer and Al-Khatib, 2001; Sousa et al., 2021; Gilbón 2023).

In addition to these primary effects, glufosinate ammonium induces a series of physiological alterations, including the generation of reactive oxygen species, reductions in photosynthetic activity, and disruption of the electron transport chain (Carbonari et al., 2016; Takano et al., 2019). These processes ultimately lead to leaf senescence and plant death.

Chlorophyll fluorescence has been widely used to assess herbicide effects on photosynthesis, especially for those that inhibit electron transport in Photosystem II (PSII) (Klem et al., 2002; Sobye et al., 2011; Dayan and de Zaccaro, 2012; Santin-Montanya et al., 2013). This technique enables the detection of early physiological effects and is also useful for herbicides that do not directly act on PSII, such as glufosinate ammonium, which nonetheless alter fluorescence patterns. Furthermore, it is an effective tool for identifying resistance mechanisms, evaluating compound penetration and detoxification, and analyzing phytotoxic interactions (Norsworthy et al., 1998). Overall, chlorophyll fluorescence is considered a sensitive biomarker for studying herbicide mode of action and activity (Dayan and de Zaccaro, 2012).

Chlorophyll fluorescence in PSII can be determined using pulse-amplitude modulation (PAM) techniques (Goltsev et al., 2016; Kalaji et al., 2018). This methodology allows for the study of PSII quantum yield during the photochemical phase (Genty et al., 1989; Baker 2008; Klughammer and Schreiber, 2008). The analysis decomposes the energy reaching PSII into different components: the photochemical process can be quantified by PSII quantum yield (Φ*_PSII_*), directly related to the electron transport rate, while the remaining energy is dissipated as heat. Part of this thermal dissipation corresponds to regulated processes (Φ*_NPQ_*), and another part to basal, non-regulated dissipation (Φ*_NO_*) (Genty et al., 1989; Kramer et al., 2004; Hendrickson et al., 2005; Klughammer and Schreiber, 2008; Chen et al., 2013; Brestic et al., 2016).

Despite its widespread use, the physiological responses of different weed species to glufosinate ammonium are not fully understood. Differential metabolism related to C3 or C4 species can influence herbicide sensitivity (Wendler et al., 1990). Some studies have reported contrasting responses between species, suggesting that factors such as herbicide absorption, translocation, and metabolic processes may play a key role in determining sensitivity (Takano et al., 2019).

Therefore, the objective of this study was to evaluate the differential physiological responses of *Lolium multiflorum* and *Echinochloa crus-galli* to glufosinate ammonium, focusing on leaf senescence, chlorophyll fluorescence dynamics, and ammonium accumulation.

## 2. Materials and Methods

### 2.1. Plant material and growth conditions

Seeds of *Lolium multiflorum* (ryegrass) glyphosate-resistant population and *Echinochloa crus-galli* (barnyardgrass) low-sensitivity population, were provided by INIA.

Seeds were germinated in Petri dishes at 25 °C in darkness. After one week, seedlings were transplanted into pots containing a 1:1 soil-to-sand mixture, with four plants per pot, later thinned to two plants per pot. Plants were watered every other day with water throughout the experiment. Growth conditions were controlled under a 12/12-hour light–dark photoperiod, with a photon flux density of 800 µmol m⁻² s⁻¹, relative humidity of 60%, and an average temperature of 25 °C.

### 2.2. Herbicide treatment

Applications were performed at the 4-leaf stage under controlled environmental conditions (25 °C, 60% RH). Glufosinate ammonium was applied at doses of 400 (D1), 500 (D2), and 600 (D3) g ha^−1^ of glufosinate ammonium (FINAL 200 SL, 200 g L⁻¹ glufosinate ammonium, Cía. Cibeles S.A.). Additionally, a control treatment (D0) without herbicide application was included. Herbicide applications were performed using a laboratory sprayer equipped with an XR8010 nozzle, calibrated to deliver 100 L ha⁻¹ at a pressure of 250 kPa.

### 2.3. Visual injury assessment

Visual damage was evaluated at different times after herbicide (glufosinate ammonium) application, according to the scale proposed by the Latin American Weed Association (ALAM, 1974). This scale establishes an index ranging from 0 to 100%, depending on the level of crop injury caused by the herbicide effect. When no visible effect is observed and the appearance is similar to the control, the index is 0%, while total plant death corresponds to 100%.

To model the visual damage data, a nonlinear regression was used, fitting a one-phase exponential decay function with a time constant parameter:

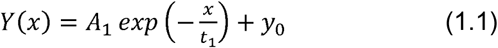

The parameters that define the model were fitted using the data from each treatment through the Damped Least Squares method, using Origin Pro 2022 software.

From the exponential decay function (1.1) determined for each treatment, the initial rate of damage progression after herbicide application was calculated. To do this, expression was derived and evaluated at x = 0, obtaining:

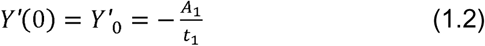

The initial rate of damage progression is an indicator of the potential level of injury caused by the herbicide at the time of application. On the other hand, the I₅₀ parameter was used as an indicator of herbicide effectiveness and speed of injury development, defined as the time required to reach 50% damage on the ALAM scale (Takano et al. 2018). This value can be determined analytically from equation (1.1) based on the model parameters, as follows:

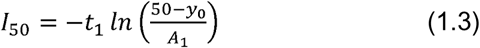

### 2.4. Energy partitioning of photosystem II: quenching and relaxation analysis

Chlorophyll fluorescence was measured through quenching and relaxation analysis, following the methodology proposed by Quero et al. (2020). Measurements were performed in vivo on the basal third of the last fully developed leaf. Readings were taken at different times after herbicide application using a PAM fluorometer (FMS1, Hansatech, Kings Lynn, UK), at 3, 6, and 9 hours after application to capture early physiological responses prior to the appearance of visible symptoms. For the quenching analysis, white actinic light at 800 µmol m⁻² s⁻¹ of photons was used. Figure 1 shows three well-defined phases during the induction of photosystem II (PSII) fluorescence (F): the initial phase, quenching analysis, and relaxation analysis

**Figure 1.**
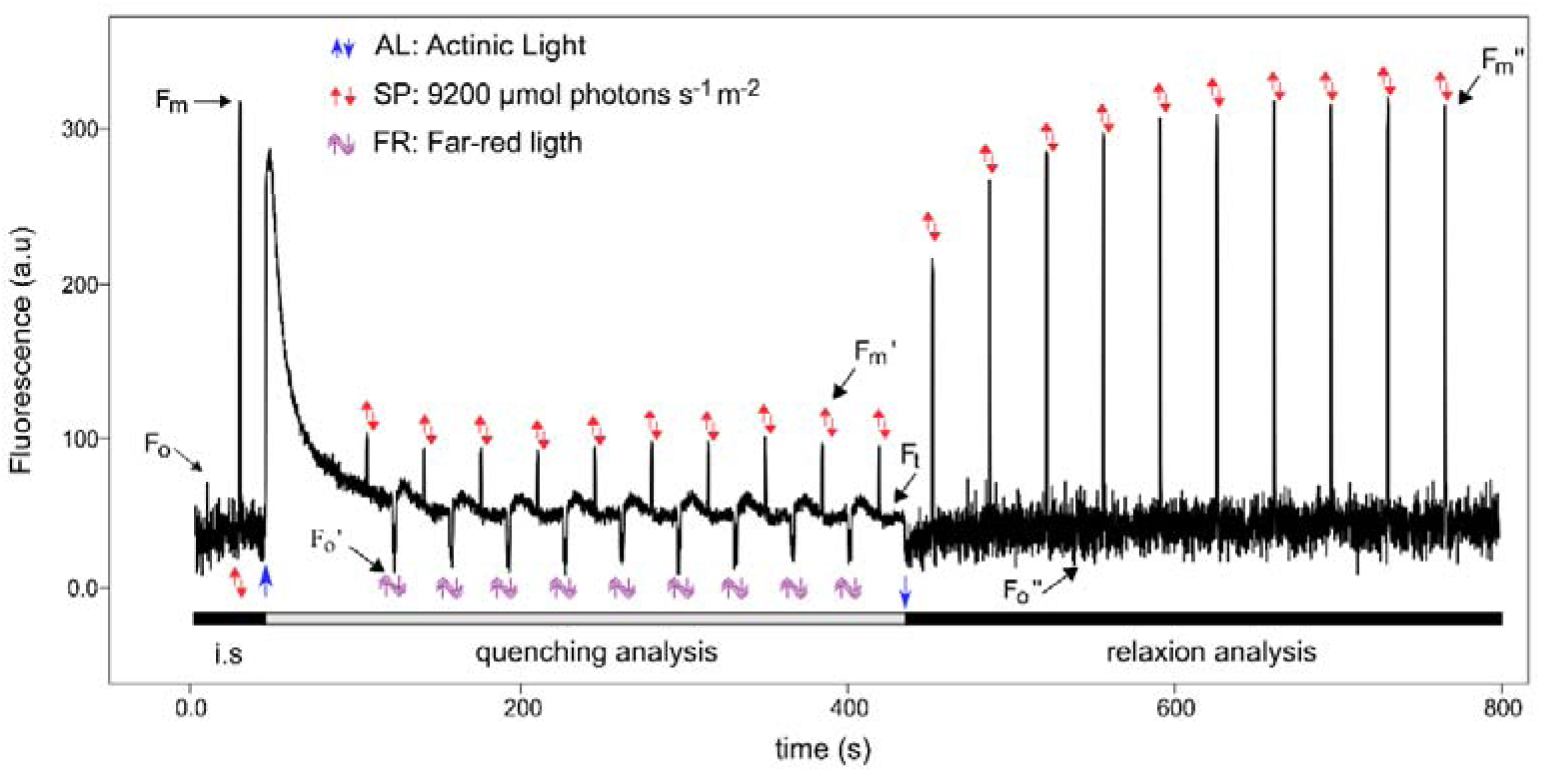
Chlorophyll fluorescence signature. Fluorescence quenching analysis followed by relaxation analysis using modulated fluorescence. The blue arrows pointing up and down indicate when the light is turned on and off, respectively. The red arrows indicate the position of the saturation pulses (SP). The double-headed arrow indicates a far-red light pulse (FR). F₀, F₀′, and F₀″ represent the minimal fluorescence in the initial phase, quenching analysis phase, and relaxation analysis phase, respectively; Fₜ is the current fluorescence for light-adapted states at time t; Fₘ and Fₘ ′ represent the maximum fluorescence under dark and light conditions, respectively. Fₘ ″ is the maximum fluorescence under dark conditions during recovery.

The quantification of energy partitioning in PSII was determined through the quantum yield of three de-excitation processes using chlorophyll fluorescence parameters (Lazár, 2015; Quero et al., 2020). This analysis is based on the premise that the sum of all de-excitation processes of the energy absorbed by PSII equals 1 (Demmig-Adams et al., 1996; Hendrickson et al., 2005; Logan et al., 2014):

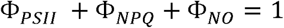

where Φ*_PSII_* is the quantum yield of PSII, Φ*_NPQ_* is the quantum yield of non-photochemical quenching, and Φ*_NO_* is the quantum yield of constitutive (basal or dark) non-regulated non-photochemical energy dissipation processes. Table 1 lists all the quantum yield parameters used in this study, along with their definitions, relationships, and references.

**Table 1.**
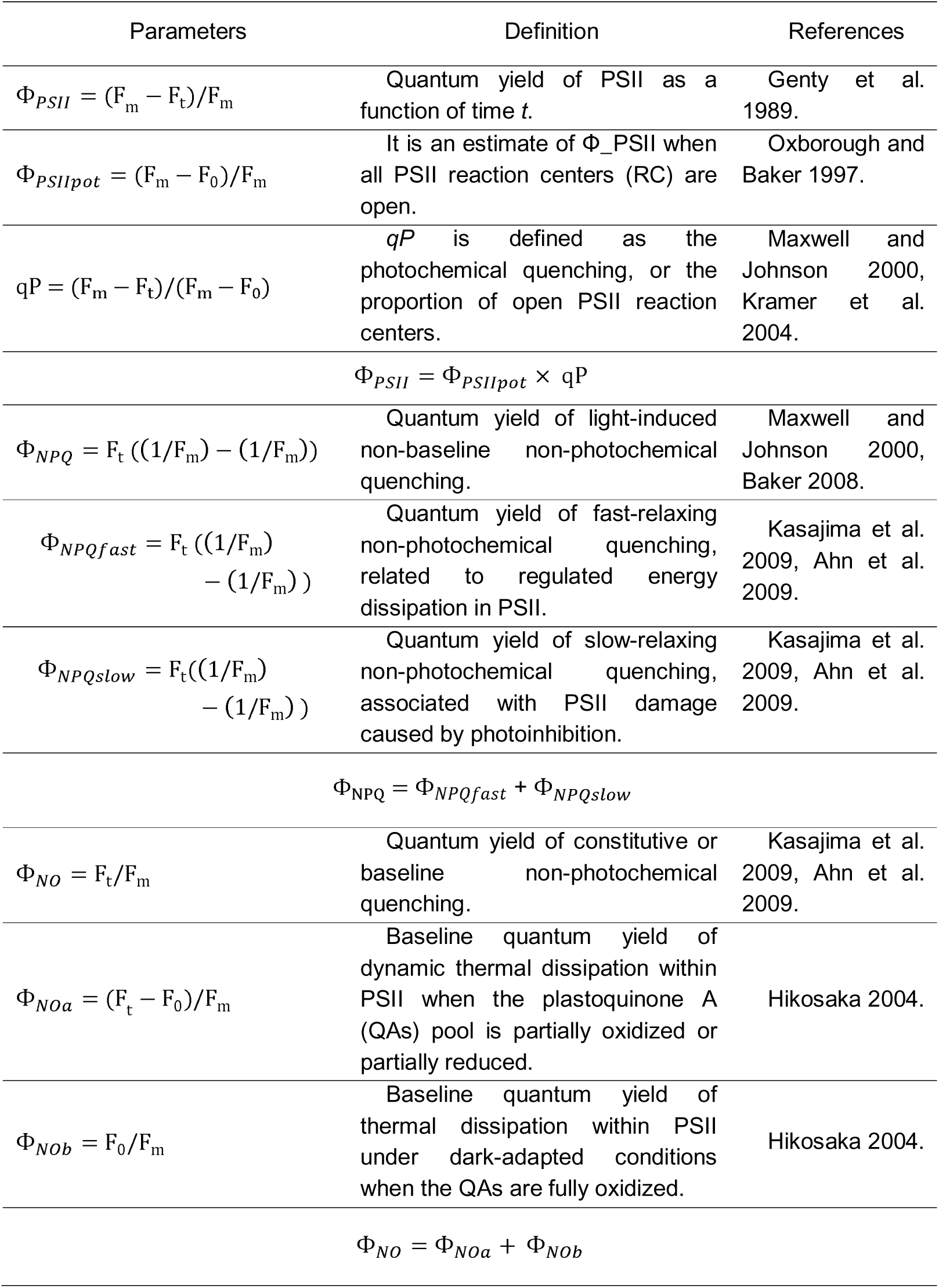
Quantum yield parameters.

| Parameters | Definition | References |
| --- | --- | --- |
| $\Phi_{PSII} = (F_m - F_t)/F_m$ | Quantum yield of PSII as a function of time $t$. | Genty et al. 1989. |
| $\Phi_{PSIIpot} = (F_m - F_0)/F_m$ | It is an estimate of $\Phi_{PSII}$ when all PSII reaction centers (RC) are open. | Oxborough and Baker 1997. |
| $qP = (F_m - F_t)/(F_m - F_0)$ | $qP$ is defined as the photochemical quenching, or the proportion of open PSII reaction centers. | Maxwell and Johnson 2000, Kramer et al. 2004. |
| $\Phi_{PSII} = \Phi_{PSIIpot} \times qP$ | | |
| $\Phi_{NPQ} = F_t((1/F_m) - (1/F_m))$ | Quantum yield of light-induced non-baseline non-photochemical quenching. | Maxwell and Johnson 2000, Baker 2008. |
| $\Phi_{NPQfast} = F_t((1/F_m) - (1/F_m))$ | Quantum yield of fast-relaxing non-photochemical quenching, related to regulated energy dissipation in PSII. | Kasajima et al. 2009, Ahn et al. 2009. |
| $\Phi_{NPQslow} = F_t((1/F_m) - (1/F_m))$ | Quantum yield of slow-relaxing non-photochemical quenching, associated with PSII damage caused by photoinhibition. | Kasajima et al. 2009, Ahn et al. 2009. |
| $\Phi_{NPQ} = \Phi_{NPQfast} + \Phi_{NPQslow}$ | | |
| $\Phi_{NO} = F_t/F_m$ | Quantum yield of constitutive or baseline non-photochemical quenching. | Kasajima et al. 2009, Ahn et al. 2009. |
| $\Phi_{NOa} = (F_t - F_0)/F_m$ | Baseline quantum yield of dynamic thermal dissipation within PSII when the plastoquinone A (QAs) pool is partially oxidized or partially reduced. | Hikosaka 2004. |
| $\Phi_{NOb} = F_0/F_m$ | Baseline quantum yield of thermal dissipation within PSII under dark-adapted conditions when the QAs are fully oxidized. | Hikosaka 2004. |
| $\Phi_{NO} = \Phi_{NOa} + \Phi_{NOb}$ | | |

### 2.5. Quantification of ammonium and nitrate

Nitrate and ammonium were extracted according to Izaguirre-Mayoral et al. (1992). A 0.2 g portion of tissue was macerated in 0.5 mL of 10.0 mM potassium phosphate buffer (pH 7.2)/96 GL ethanol (1:1). The homogenate was centrifuged at 5000 × g for 10 minutes at room temperature. The supernatant was used for the quantification of nitrate and ammonium.

Nitrate was quantified according to Cataldo et al. (1975) with modifications. To 0.1 mL of sample, 0.4 mL of 0.36 M salicylic acid in 18.4 M sulfuric acid was added and mixed. After 20 minutes, 5.0 mL of 3.8 M NaOH was added. Absorbance was measured at 410 nm, using KNO₃ as the standard.

Ammonium was quantified following Solorzano (1969). To 1.0 mL of sample, diluted to an appropriate concentration, the following reagents were added in order:

- 0.2 mL of 1.1 M phenol (10% w/v in absolute ethanol)
- 0.2 mL of 16.8 mM sodium nitroprusside (0.5% w/v in water)
- 0.4 mL of 0.68 M sodium citrate (20% w/v) – 0.3 M NaOH (1% w/v in water)
- 0.1 mL of 1.3 M NaClO (10%)

After each addition, the mixture was stirred and then incubated in darkness for 2 hours. Absorbance was measured at 640 nm, using NH₄Cl as the standard.

### 2.6. Statistical analysis

The experiment was conducted using a completely randomized design with four replicates per treatment. The experimental unit consisted of a pot containing two plants. Treatments were defined by the interaction among the levels of the genotype factor (*E. crus-galli* and *L. multiflorum*), the herbicide factor (D0: 0 g ha⁻¹ (control), D1: 400 g ha⁻¹ glufosinate ammonium, D2: 500 g ha⁻¹ glufosinate ammonium, and D3: 600 g ha⁻¹ glufosinate ammonium), and the time factor (3, 6, and 9 hours after application).

For statistical analysis, the following general linear factorial model was used:

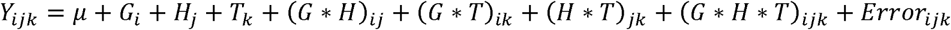

*G_i_* is the effect of the i-th genotype (*E. crus-galli* and *L. multiflorum*); *H_j_* is the effect of the j-th herbicide level (D0: 0 g ha⁻¹—control—; D1: 400 g ha⁻¹ glufosinate ammonium; D2: 500 g ha⁻¹ glufosinate ammonium; and D3: 600 g ha⁻¹ glufosinate ammonium); and *T_k_* is the effect of the k-th time (3, 6, and 9 hours after herbicide pplication). (*G* ∗ *H*)*_ij_* represents the interaction between the i-th genotype and the j-th herbicide level; (*G* ∗ *T*)*_ik_* represents the interaction between the i-th genotype and the k-th time; (*H* ∗ *T*)*_jk_* represents the interaction between the j-th herbicide level and the k-th time; and (*G* ∗ *H* ∗ *T*)*_ijk_* represents the interaction among the i-th genotype, the j-th herbicide level, and the k-th time. *Error_ijk_* is the random error associated with each experimental unit.

The model used for the statistical analysis of nitrate and ammonium accumulation was the same general linear model with a factorial arrangement used for chlorophyll fluorescence, excluding the *T_k_* term since measurements were taken only 9 hours after glufosinate-ammonium application.

After verifying the assumptions of normality (Shapiro–Wilk test) and homogeneity of variance (Levene’s test), an analysis of variance (ANOVA) was performed for each variable to determine whether there were significant effects of the factors. Differences between means were assessed by orthogonal contrast analysis (p ≤ 0.05). All statistical analyses were performed in R using the stats package (R Core Team, 2023). The model was fitted using the lme4 package (Bates et al., 2015). The best linear unbiased estimators (BLUEs) and contrast analyses were obtained using the emmeans package (Lenth, 2025).

## 3. Results and Discussion

### 3.1. Visual injury

In Figure 2 (a) and (b), the progression of visual injury in both species following glufosinate ammonium application at different doses is shown. No visual symptoms were observed within the first 48 hours after application. Takano et al. (2019) reported the same situation in studies conducted on ryegrass. That author indicated that, at a dose of 560 g ha⁻¹ glufosinate ammonium (intermediate between D2 and D3), the first visual symptoms begin to appear around 48 hours after application. Likewise, the absence of visual symptoms does not indicate the absence of leaf senescence processes.

**Figure 2.**
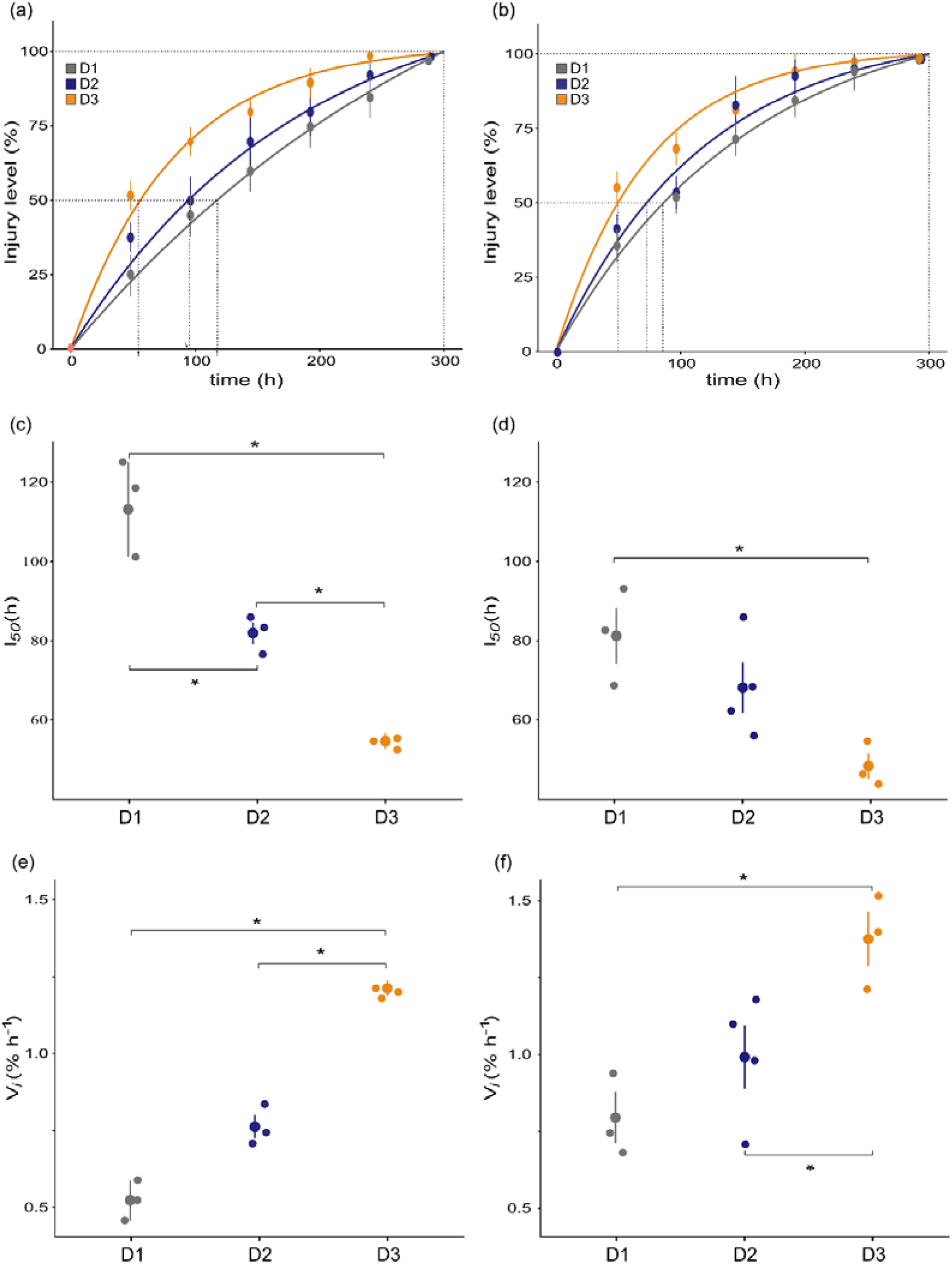
Response to glufosinate application. (a), (b) Graph of injury level (%) as a function of time (hours) after glufosinate ammonium application; (c), (d) Graph of I₅₀ as a function of herbicide dose; and (e), (f) Graph of initial injury progression rate (% day⁻¹) for (a), (c), and (e) *Lolium multiflorum* (L.) and (b), (d), and (f) *Echinochloa crus-gall*i (barnyardgrass). D1: 400 g ha⁻¹ glufosinate ammonium; D2: 500 g ha⁻¹ glufosinate ammonium; D3: 600 g ha⁻¹ glufosinate ammonium.

It is important to note that, when evaluating each species, different *I₅₀* values were obtained depending on the herbicide dose used. In both species, the *I₅₀* parameter was lower in treatment D3 compared to D2 and D1.

However, differences between D2 and D3 were observed only in ryegrass (Figure 2 (c) and (d).

When comparing between species, it can be observed that ryegrass plants exposed to treatment D1 (lowest herbicide dose) needed more than 100 hours to reach *I₅₀* (Figure 2c). In contrast, barnyardgrass plants exposed to all three doses reached *I₅₀* in less than 100 hours (Figure 2d). *I₅₀* values were 113 and 81 hours after application in treatment D1 for ryegrass and barnyardgrass, respectively. The results obtained for the level of injury in ryegrass following glufosinate ammonium application are consistent with those reported by Takano et al. (2019).

In Figure 2 (e) and (f), the initial rate of injury progression is shown. In both species, it can be observed that the rate of symptom expression depends on the herbicide dose (D1, D2, and D3).

Responses varied among species, with slower damage progression in ryegrass. At dose 1 (D1), ryegrass showed 0.52% damage/hour (Figure 2e) and barnyardgrass showed 0.79% damage/hour (Figure 2f), while at dose 3 (D3), ryegrass showed 1.21% damage/hour (Figure 2e) and barnyardgrass showed 1.38% damage/hour (Figure 2f). This result is consistent with that reported by Takano et al. (2019), who found that ryegrass exhibits lower sensitivity to glufosinate ammonium compared to other C4 weed species evaluated.

A species-dependent response was observed; the lower sensitivity of ryegrass was consistent with results from Takano et al. (2019) and Culpepper et al. (2000), who also reported lower sensitivity to glufosinate ammonium in C3 species.

### 3.2. Chlorophyll fluorescence of photosystem II

Chlorophyll fluorescence kinetics were evaluated in ryegrass and barnyardgrass under the same experimental conditions used for visual injury assessment. Although glufosinate primarily affects nitrogen metabolism, its indirect effects on photosynthesis make fluorescence analysis a valuable tool for assessing early physiological responses. From the first evaluation time (48 h) and throughout the experiment, the chlorophyll fluorescence kinetics did not maintain their integrity (Supplementary Material Figure S1), making the assessment of energy partitioning highly complex.

Considering this result and the observed visual damage, we decided to shorten the interval between herbicide application and evaluation to measure chlorophyll fluorescence before visual damage became evident.

Different herbicide doses caused alterations in the electron transport chain at 6 and 9 hours after application (Figures 3 and 4). These measurements were useful for evaluating energy distribution within photosystem II and determining the fate of the energy absorbed during photosynthesis. In contrast, the chlorophyll fluorescence kinetics of ryegrass showed no detectable changes across doses or sampling times. Barnyardgrass (Figure 3) exhibited more pronounced photosystem II damage than ryegrass (Figure 4), indicating that the two species differ in their photosystem II sensitivity.

**Figure 3.**
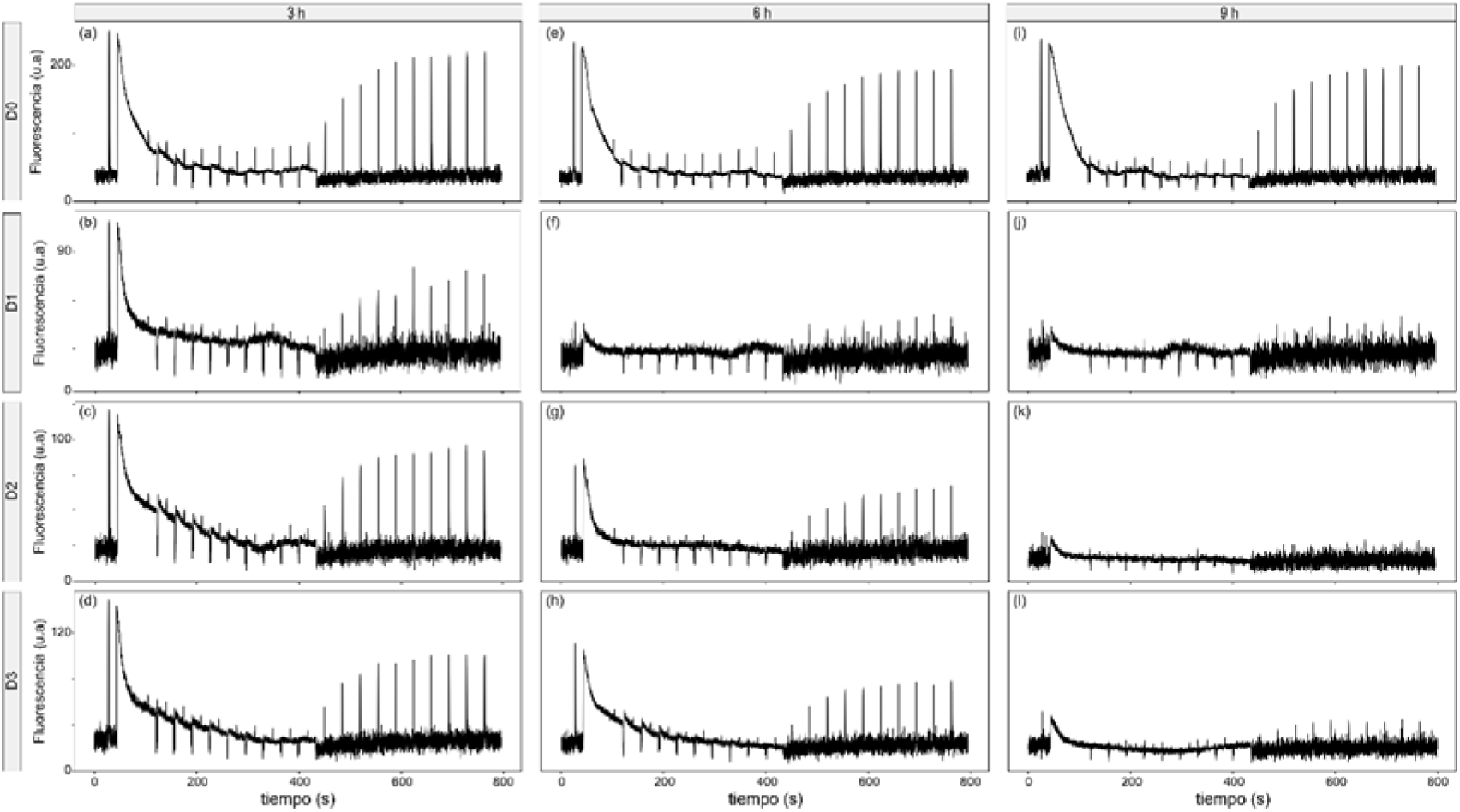
Chlorophyll fluorescence traces of photosystem II in *Echinochloa crus-galli* (barnyardgrass). under four glufosinate ammonium doses at three time points after treatment application (3, 6, and 9 hours). (a), (e), (i) D0: 0 g L ha⁻¹ (control); (b), (f), (j) D1: 400 g ha⁻¹ glufosinate ammonium; (c), (g), (k) D2: 500 g ha⁻¹ glufosinate ammonium; (d), (h), (l) D3: 600 g ha⁻¹ glufosinate ammonium.

**Figure 4.**
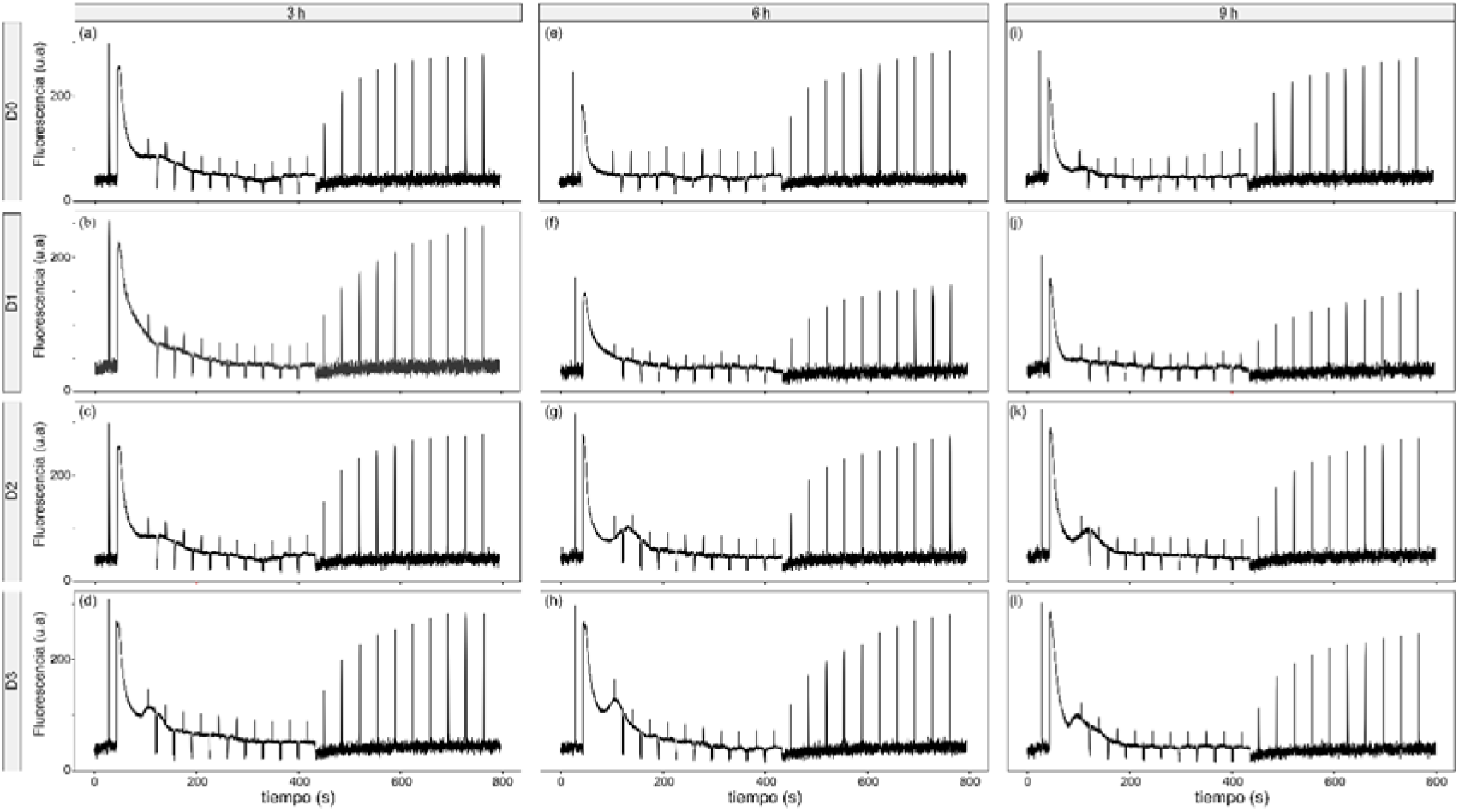
Chlorophyll fluorescence traces of photosystem II in *Lolium multiflorum* (ryegrass) under four doses of glufosinate ammonium at three time points after treatment application (3, 6, and 9 hours). (a), (e), (i) D0: 0 g ha⁻¹ (control). (b), (f), (j) D1: 400 g ha⁻¹glufosinate ammonium. (c), (g), (k) D2: 500 g ha⁻¹ glufosinate ammonium. (d), (h), (l) D3: 600 g ha⁻¹ glufosinate ammonium.

Sensitivity to glufosinate-ammonium has been previously described as genotype-dependent. Some studies found that inhibition was slower in C4 plants (Wendler et al., 1990). Our results, where ryegrass showed the greatest tolerance, are consistent with those reported by Takano et al. (2019).

Differences in sensitivity to glufosinate ammonium are related to differences in translocation (Kumaratilake et al., 2002). Structural differences may also exist, influencing absorption and translocation and affecting herbicide sensitivity (Galván et al., 2010).

### 3.3. Energy partitioning in photosystem II

The analysis of variance (ANOVA) at the three evaluation times (3, 6, and 9 hours after herbicide application) showed that none a significant triple interaction among genotype, herbicide dose, and time (Supplementary Material Table S1). Therefore, subsequent analyses were conducted of the main factors and their two-way interactions.

Based on the ANOVA results, the main factor species was significant for all three parameters. In addition, for the parameter Φ*_PSII_* both the herbicide dose factor and the genotype × herbicide interaction was significant. The parameter Φ*_PSII_* was not affected by time after application, indicating that it was similarly influenced at 3, 6, and 9 hours after herbicide application.

For the parameter Φ*_NO_* all factors and their interactions were significant. This parameter represents energy dissipated through basal (non-photochemical) processes. Unlike Φ*_PSII_* (the actual efficiency of photosystem II), Φ*_NO_* was affected by time.

Φ*_NO_* is associated with genes encoding structural proteins of PSII, suggesting that non-photochemical energy dissipation processes are related to the structural composition of PSII (Quero et al., 2020). Moreover, Φ*_NO_* has been described as an indicator of PSII damage (Klughammer and Schreiber, 2008). In our case, Φ*_NO_* reflects the effect of elapsed time since glufosinate-ammonium application, indicating structural damage in photosystem II.

Regarding the parameter Φ*_NPQ_*, only the main factor genotype showed a significant effect, while the other factors and their interactions were not significant.

In barnyardgrass, treatment control, the plant allocated 40% of the received energy to the photochemical phase of photosynthesis (Φ*_PSII_*), while the remaining energy was dissipated as heat—40% through non-basal processes (Φ*_NPQ_*), and 20% through basal dissipative processes Φ*_NO_*. In treatments with herbicide reduced the proportion of energy allocated to photochemical processes and increased energy dissipation as heat. This shift indicates a disruption of electron transport and reduced photosynthetic efficiency following herbicide application. That result is consistent with reported by Jeong et al. (2024) reporting reductions in PSII efficiency.

Regarding Φ*_NPQ_*, no significant differences were observed across the doses and time points evaluated (Figure 5a). As for Φ*_NO_*, the range of energy allocated to basal dissipative processes varied between 20% and 60% (Figure 5a). At 3 hours after glufosinate-ammonium application, treatments D1 and D2 increased the value of Φ*_NO_* to 30%, differing from treatment D0, which was 20%. At 6 hours after glufosinate ammonium application, treatments D1 and D2 continued to increase the value of Φ*_NO_* to 60% and 40%, respectively, differing from treatment D0, which was 20% (Figure 5a). At 9 hours, D1, D2, and D3 dissipated more heat through basal processes (50%, 40%, and 40%, respectively) compared to D0 (20%). This result indicates a greater proportion of energy allocated to basal energy dissipation processes in plants treated with glufosinate ammonium, with this effect magnifying over time after application. It has been reported that glufosinate has the potential to limit its own translocation due to rapid cellular damage (Beriault et al., 1999), which could explain why higher doses of glufosinate-ammonium showed a delayed effect on the evaluated variables compared to lower doses.

**Figure 5.**
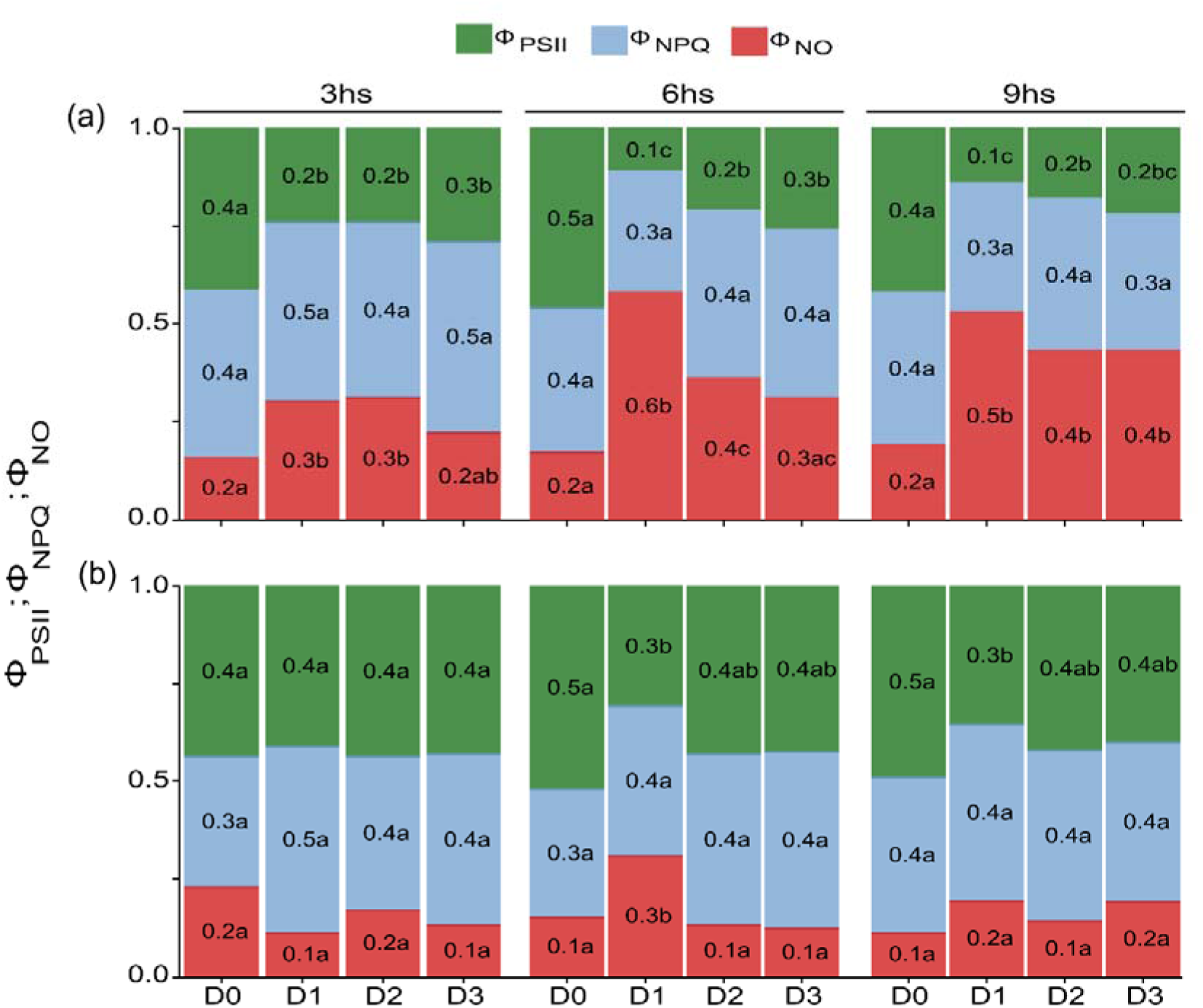
Energy partitioning parameters of photosystem II. (a) Echinochloa crus-galli (L.) Beauv. and (b) Lolium multiflorum (L.) under four doses of glufosinate ammonium at three times after treatment application (3, 6, and 9 hours). D0: 0 g ha⁻¹ (control). D1: 400 g ha⁻¹ glufosinate ammonium. D2: 500 g ha⁻¹ glufosinate ammonium. D3: 600 g ha⁻¹glufosinate ammonium.

In barnyardgrass, it can be concluded that, after glufosinate ammonium application, there was a decrease in Φ*_PSII_*, caused by a deficiency in electron transfer to the photosynthetic photochemical process (Murata et al., 2007; Muh et al., 2012). In turn, the observed increase in Φ*_NO_* indicates the presence of non-regulated heat dissipation, which serves as an indicator of damage to the photosystem structure (Klughammer and Schreiber, 2008). Moustaka and Moustakas (2014) report decreases in the quantum yield of photosystem II (Φ*_PSII_*), following applications of Paraquat (a herbicide that interferes with electron transport), while they observed increases in Φ*_NPQ_* and decreases in Φ*_NO_*, these results are associated with a rapid photoprotective mechanism. In the case of Paraquat, the excess energy dissipated (high Φ*_NPQ_*) causes a decrease in ETR and explains the mode of action of Paraquat. In our case, our results are consistent with Moustaka and Moustakas (2014) regarding the effect of Φ*_PSII_*, but it does not affect the other two parameters evaluated. In barnyardgrass, the increases in Φ*_NO_* are not related to a photoprotective mechanism, but rather to the disruption of the photosystem structure. The observed effect on the parameter Φ*_NO_* in barnyardgrass (increase of Φ*_NO_*) is consistent with the results reported by Zhang et al. (2022), when wheat was grown under nitrogen-deficient conditions.

The ryegrass control plants (D0) allocated 50% of the absorbed energy to the photochemical phase of photosynthesis, while the remainder was dissipated as heat: 30–40% to non-basal processes and 10–20% to basal dissipative processes (Figure 5a). It was observed that the energy distribution in both species in control was similar. While barnyardgrass exhibits an effect of herbicide by increasing its Φ*_NO_*, ryegrass does not.

In ryegrass, herbicide-treated plants (D1, D2, and D3) showed energy distribution ranges similar to the control D0. At the evaluation times, this genotype did not exhibit clear increases or decreases in the measured parameters. These results are consistent with those obtained in the visual damage analysis, as both aspects of senescence are delayed in ryegrass compared to barnyardgrass. These findings agree with those reported by Takano et al. (2019), where ryegrass shows lower sensitivity to glufosinate ammonium than other evaluated C4 weeds.

Furthermore, this analysis allows us to decompose some principal factors to continue investigating what occurs between the genotypes. The parameter Φ*_NO_* consists of Φ*_NOb_*, which is related to heat dissipation through the reaction centers, and Φ*_NOa_*, associated with the antenna complex (Quero et al., 2020). Φ*_NPQ_* is composed of non-photochemical quantum yields of rapid relaxation (Φ*_NPQfast_*) and slow (Φ*_NPQslow_*) (Kasajima et al. 2009), while Φ*_NPQfast_* is related to regulated heat dissipation in PSII and Φ*_NPQslow_* is related to the damage caused by photoinhibition (Ahn et al., 2009; Kasajima et al., 2009).

Regarding barnyardgrass (Figure 6a), it can be observed at the three evaluated time points (3, 6, and 9 hours) that D0 dissipates between 6% and 9% of energy through the reaction centers(Φ*_NOb_*), while 10% is allocated to heat dissipation via the antenna complex (Φ*_NOa_*). It can be inferred that in control treatments, the proportion of energy dissipated through the reaction centers and the antenna complex is similar.

**Figure 6.**
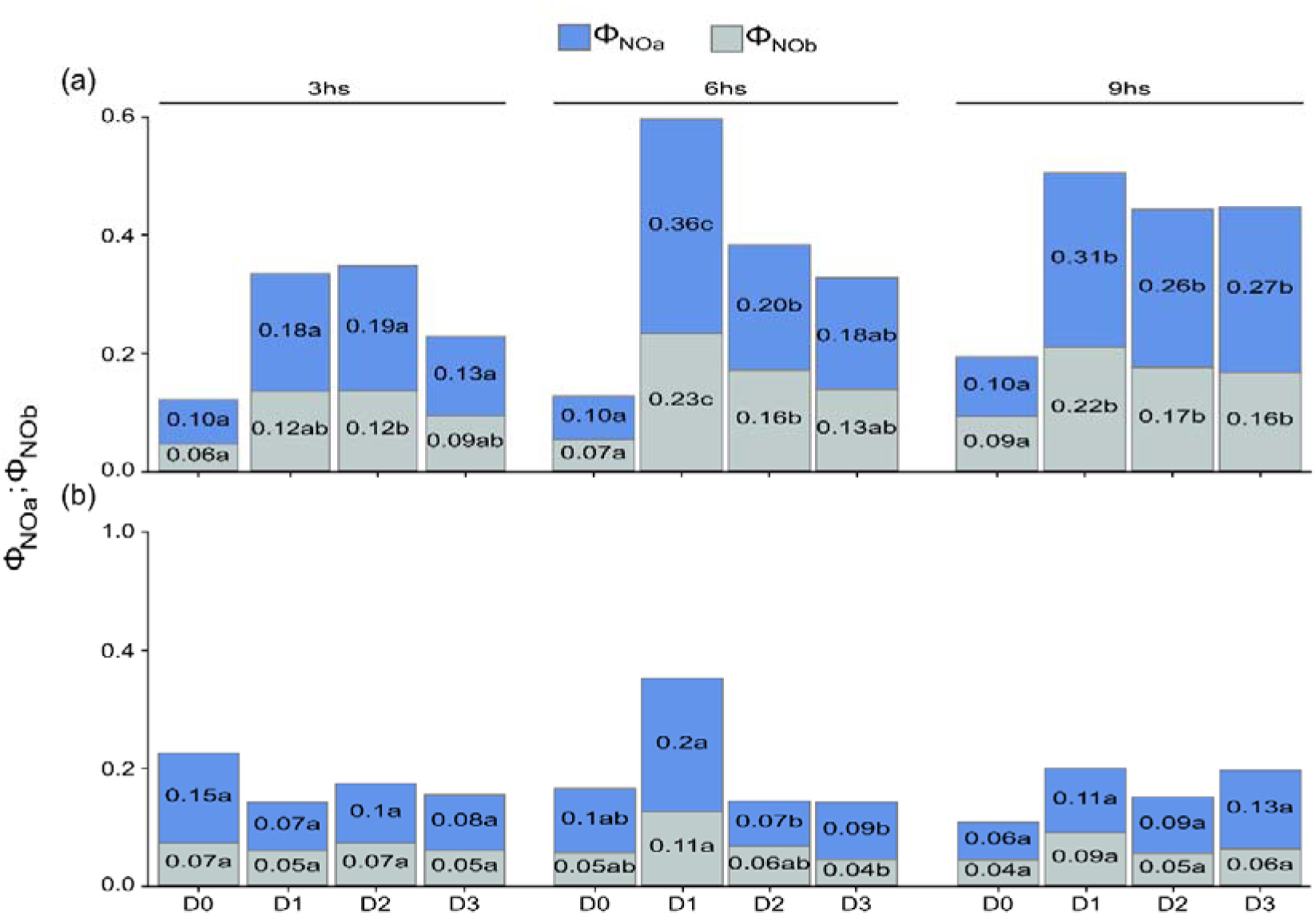
Sub-parameters Φ*_NOa_* y Φ*_NOb_* of basal energy dissipation from non-regulated processes. (a) *Echinochloa crus-galli* (L.) Beauv. and (b) *Lolium multiflorum* (L.) under four doses of glufosinate ammonium at three time points after treatment application (3, 6, and 9 hours). D0: 0 g ha⁻¹ (control). D1: 400 g ha⁻¹ glufosinate ammonium. D2: 500 g ha⁻¹glufosinate ammonium. D3: 600 g ha⁻¹ glufosinate ammonium.

When observing treatments D1, D2, and D3 in barnyardgrass at the three evaluation times, these dissipate between 9% and 23% of energy through the reaction centers, while 13% to 36% is dissipated via the antenna complex (Figure 6a). Observing Figure 6 (a), it can be inferred that as time progresses after herbicide application, in barnyardgrass the energy dissipation through both the antenna complex and the reaction centers increases, differentiating from the control. Furthermore, it can be inferred that the increase in Φ*_NO_* is determinated by Φ*_NOa_*, indicating that the greatest energy dissipation is occurring through the antenna complex.

Observing the response in ryegrass (Figure 6b), it can be seen at the three evaluated time points (3, 6, and 9 hours) that D0 dissipates between 4% and 7% of energy through the reaction centers, while 6% to 15% is allocated to heat dissipation via the antenna complex. In treatments D1, D2, and D3 at the three evaluation times, energy dissipation through the reaction centers ranged from 5% to 11%, while 7% to 20% was dissipated via the antenna complex.

In ryegrass, herbicide-treated plants (D1, D2, and D3) showed no differences in basal energy dissipation from non-regulated processes compared to control D0 (Figure 6b). In barnyardgrass, the parameters Φ*_NO_* y Φ*_NOa_* are higher than in ryegrass, indicating damage to the structure of photosystem II. These results are consistent with those obtained in the visual damage analysis, as both aspects of senescence are delayed in ryegrass compared to barnyardgrass. These findings agree with those reported by Takano et al. (2019), where ryegrass shows lower sensitivity to glufosinate ammonium than the evaluated C4 weeds.

As shown in Figure 7, within each genotype no significant differences were observed in the evaluated parameters (Φ*_NPQfast_* and Φ*_NPQslow_*)) among treatments D0, D1, D2, and D3. In Figure 7(a), barnyardgrass exhibited values ranging from 23 to 39% for regulated heat dissipation (Φ*_NPQfast_*) and from 2 to 12% for heat dissipation associated with photodamage (Φ*_NPQslow_*, regardless of the herbicide dose. In contrast, ryegrass showed regulated thermal dissipation values between 32 and 43%, while losses due to photodamage were nearly negligible.

**Figure 7.**
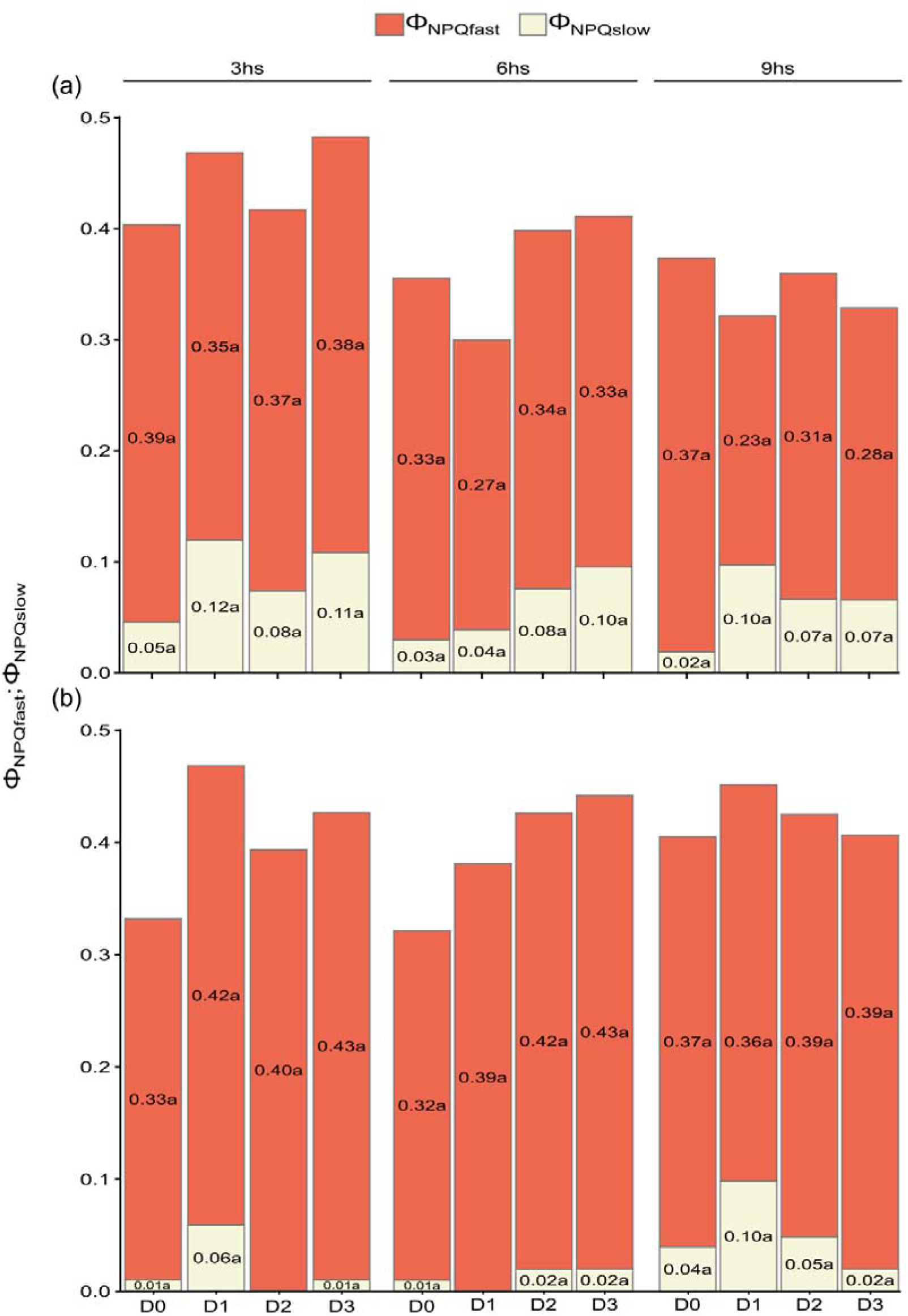
Sub-parameters Φ*_NPQfast_* y Φ*_NPQslow_* of non-photochemical heat dissipation through regulated processes. (a) *Echinochloa crus-galli* (L.) Beauv. and (b) *Lolium multiflorum* (L.) under four doses of glufosinate-ammonium at three time points after treatment application (3, 6, and 9 hours). D0: 0 g ha⁻¹ (control). D1: 400 g ha⁻¹ glufosinate-ammonium. D2: 500 g ha⁻¹ glufosinate-ammonium. D3: 600 g ha⁻¹ glufosinate-ammonium.

For both genotypes, we can identify that Φ*_NPQfast_* has greater relevance in Φ_NPQ due to its proportion. This result is consistent with that reported by Shrestha et al. (2012) under low nitrogen supply conditions in rice plants. According to Shrestha et al. (2012), reduced nitrogen availability induces xanthophyll cycle activity, increasing Φ*_NPQfast_* . Φ*_NPQslow_* shows no significance between doses and post-application times of the herbicide, indicating that the herbicide dose has no effect on photoinhibition.

To complement the analysis of the main parameters of light partitioning, the following parameters were evaluated: Φ*_PSIIpot_* which represents the maximum quantum yield in light and *qP* which determines the proportion of open reaction centers at that moment. Φ*_PSIIpot_* describes the stability and maximum capacity of PSII, whereas *qP* explains the cause of the variations in quantum yield (Kramer et al., 2004; Kasajima et al., 2009; Lazár, 2015).

In Figure 8, for barnyardgrass, treatment D0 showed a potential quantum light yield of PSII (Φ*_PSIIpot_*) ranging from 72% to 79%, which was higher than the values observed in the herbicide-treated treatments (D1, D2, and D3), which ranged from 65% to 71%. Regarding the proportion of open reaction centers (*qP*), a more pronounced difference was observed between treatment D0, which ranged from 53% to 57%, and treatments D1, D2, and D3, which ranged from 17% to 34% (Figure 8 a-c).

**Figure 8.**
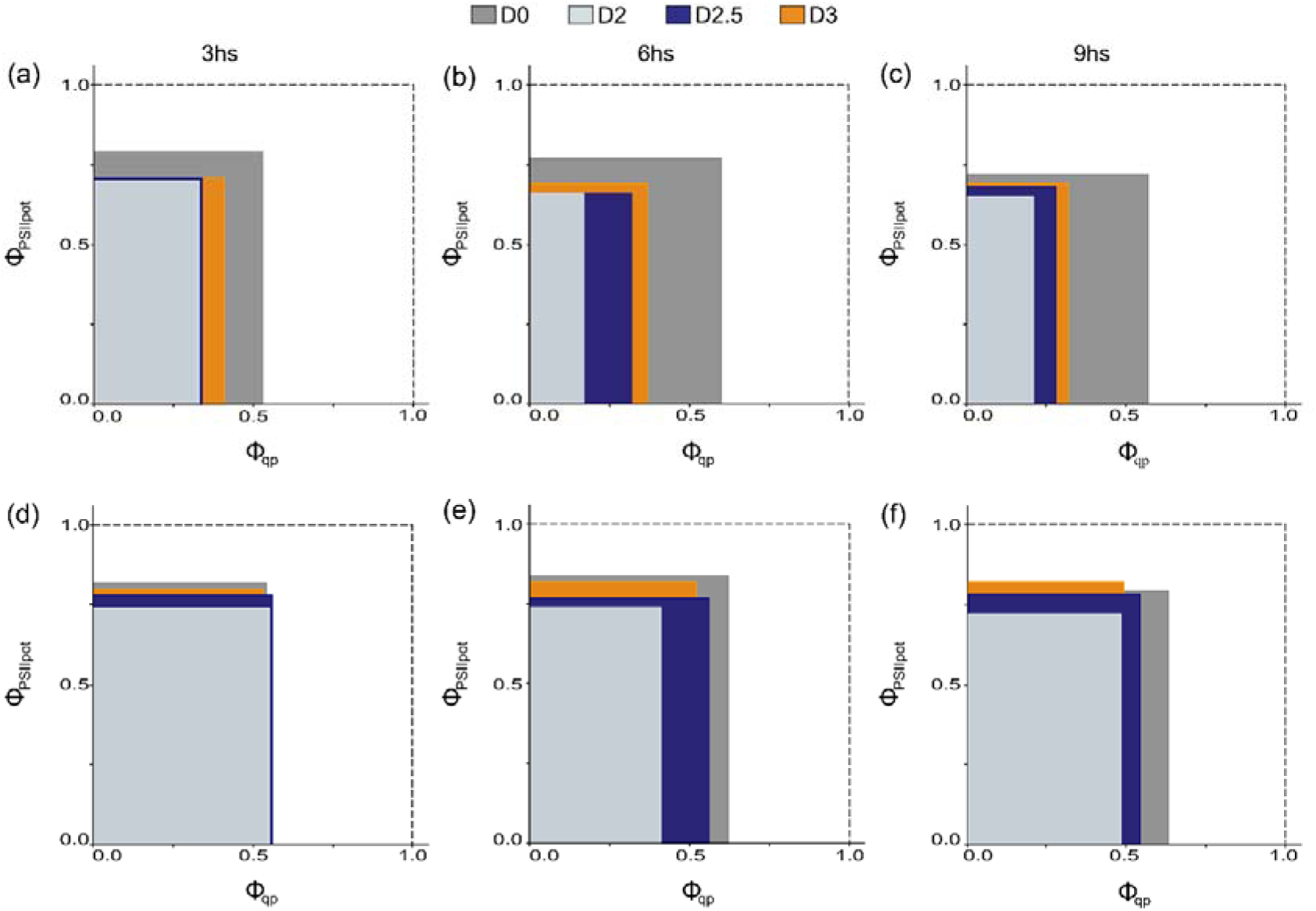
Quantification of the maximum quantum yield in light (Φ*_PSIIpot_*) and the proportion of open reaction centers (*qP*) a, b, and c. Echinochloa crus-galli (L.) Beauv. after 3, 6, and 9 hours of treatment application, respectively d, e, and f. Lolium multiflorum (L.) after 3, 6, and 9 hours of treatment application. D0: 0 g ha⁻¹ (control). D1: 400 g ha⁻¹ glufosinate ammonium. D2: 500 g ha⁻¹ glufosinate ammonium. D3: 600 g ha⁻¹ glufosinate ammonium.

**Figure 9.**
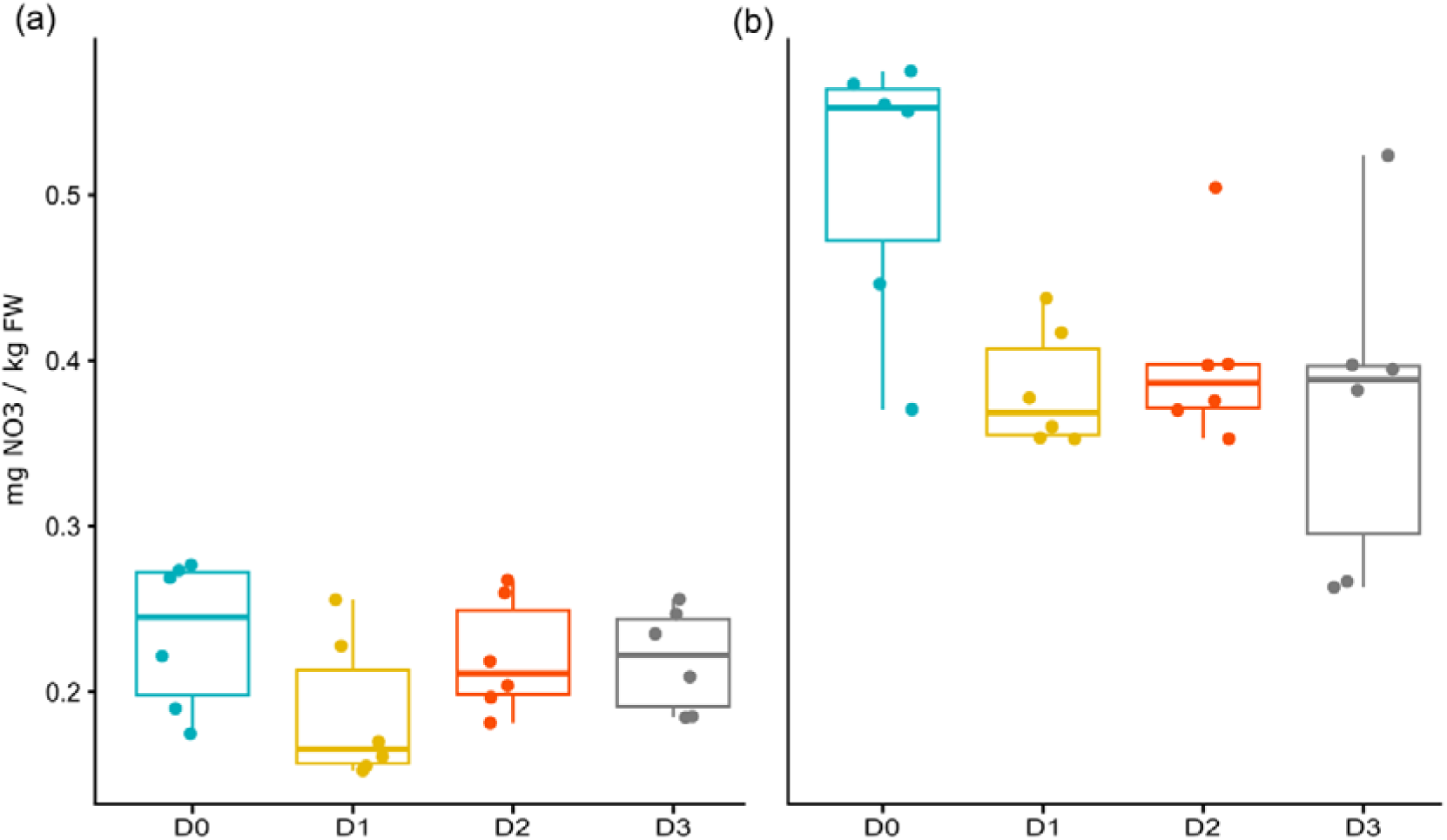
Nitrate quantification (mg NO₃/kg FW) (a) Echinochloa crus-galli (L.) Beauv. and (b) Lolium multiflorum (L.) 9 hours after treatment application. D0: 0 g ha⁻¹ (control). D1: 400 g ha⁻¹ glufosinate ammonium. D2: 500 g ha⁻¹ glufosinate ammonium. D3: 600 g ha⁻¹ glufosinate ammonium.

For this genotype, there is a clear effect of time on the parameter *qP*, as time increases after application, this parameter decreases in treatments D1, D2, and D3. A greater impact on the parameter *qP* indicates that the reaction centers begin to close starting 3 hours after glufosinate ammonium application, and this effect becomes more pronounced with increasing time after application, this explains the decrease in Φ*_PSII_*. This effect is consistent with that observed by Moustaka and Moustakas (2014), where following a Paraquat application (a herbicide that interferes with electron transport), *qP* it decreased by 13.5% one hour after herbicide application.

Regarding the ryegrass genotype, no clear effect was observed in the evaluated parameters. Φ*_PSIIpot_* takes values from 72 to 84% across the four treatments, and ranges from 62 to 54% (Figure 8 d-f). When comparing the impact on these parameters in each species, control efficiencies and reduced sensitivity were observed starting 3 hours after glufosinate ammonium application. These results are consistent with those obtained in the visual damage analysis and the main fluorescence parameters, where senescence in ryegrass was delayed compared to barnyardgrass.

Observing the effects on the parameters *qP* and Φ*_NOb_*, both related to the reaction center of photosystem II, in the barnyardgrass genotype, glufosinate ammonium-treated plants differ from the control treatment without herbicide. Moreover, this effect becomes more pronounced over time. The effect on both parameters indicates that, starting 3 hours after herbicide application, the reaction centers begin to close, and as time progresses, there is a trend toward decreased energy dissipation through the reaction centers. This suggests a dependence between the opening of the reaction centers and the structural composition of the PSII reaction centers. As described by Fleitas and Pereyra (2022), the structural conformation of the reaction centers Φ*_NOb_* conditions the opening of the reaction centers *qP*.

Overall, the evaluated photosynthetic parameters revealed rapid physiological changes following glufosinate ammonium application, allowing early detection of herbicide effects before visible symptoms appear. These results highlight the potential of chlorophyll fluorescence as a sensitive and reliable tool for assessing herbicide activity and plant responses.

### 3.3. Quantification of nitrate and ammonium

Nitrate concentration was not significantly affected by species, herbicide dose, or their interaction.

Nitrate levels remained similar across treatments in both species, with no significant differences between control (D0) and herbicide-treated plants.

In contrast, of ammonium quantification, we determined that the variables species and the species-glufosinate interaction are significant, indicating that ammonium accumulation varies depending on the species and the applied dose.

In Figure 10, it can be observed that in both barnyardgrass and ryegrass, treatment D0 presented ammonium values close to 0. In barnyardgrass, ammonium accumulation for D1, D2, and D3 ranged between 0.159 and 0.456 mg NH₃/kg FW, while in ryegrass, ammonium quantification for the same treatments ranged between 0.131 and 0.631 mg NH₃/kg FW. In neither species there was a statistically significant difference in ammonium accumulation among D1, D2, and D3.

**Figure 10.**
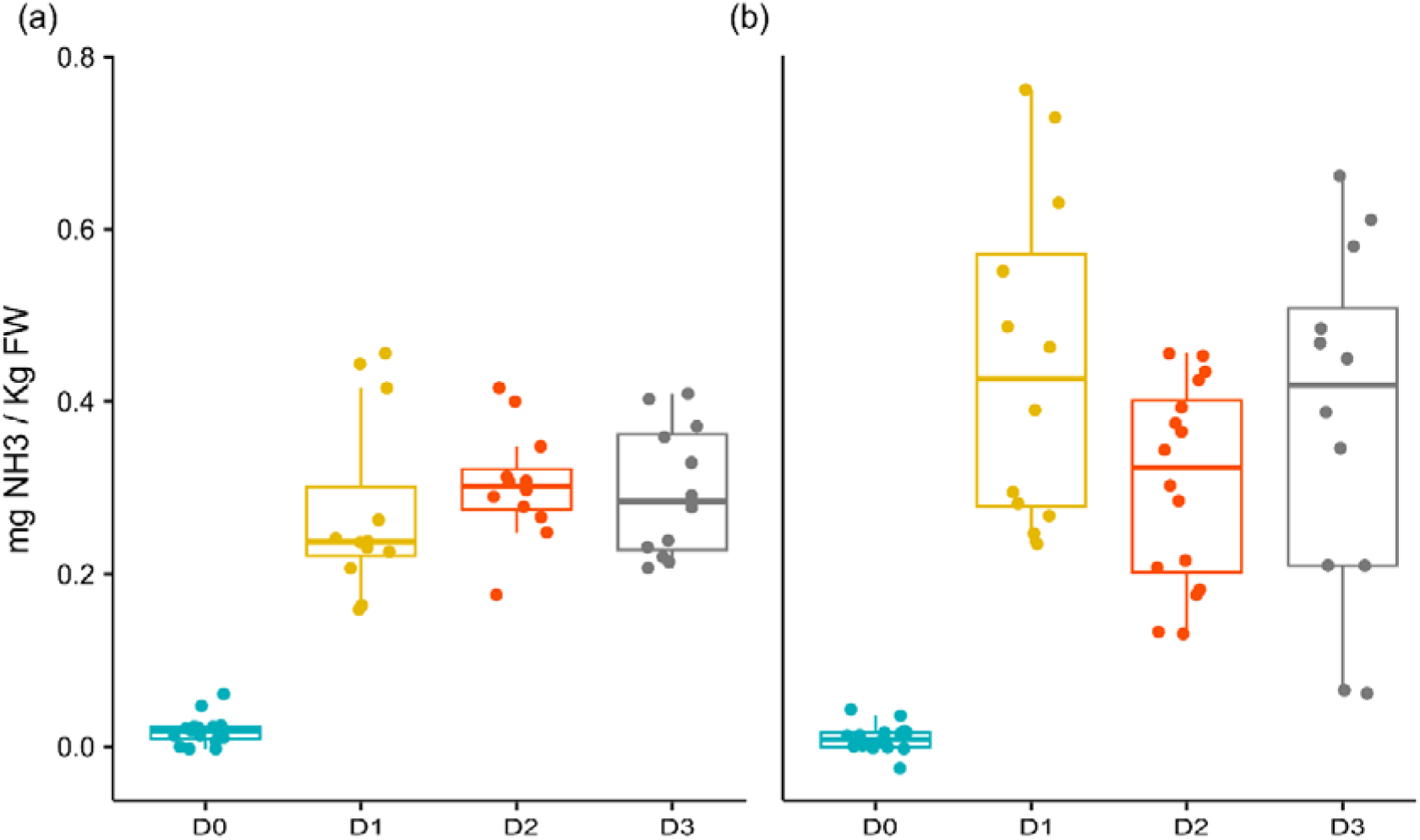
Ammonium quantification (mg NH₃/kg FW) (a) *Echinochloa crus-galli* (L.) Beauv. and (b) *Lolium multiflorum* (L.) 9 hours after treatment application. D0: 0 g ha⁻¹ (control). D1: 400 g ha⁻¹ glufosinate ammonium. D2: 500 g ha⁻¹ glufosinate ammonium. D3: 600 g ha⁻¹ glufosinate ammonium.

The effect of glufosinate ammonium observed in different doses compared to D0 on ammonium accumulation is evident and can be explained by the mode of action of this herbicide. Inhibition of glutamine synthetase activity leads to rapid ammonium accumulation (Wendler et al., 1990; Chompoo and Pornprom, 2008; Puertas de Freitas e Silva et al., 2016; Takano et al., 2019), which causes cell destruction (Senseman, 2007). Therefore, ammonium increase is used as an indicator of glufosinate ammonium efficacy (Pornprom et al., 2000; Petersen and Hurle, 2001).

Barnyardgrass accumulated lower ammonium levels than ryegrass (Figure 10 a and b), which is consistent with previous reports indicating reduced ammonium accumulation in C4 species due to lower photorespiration rates (Wendler et al., 1990).

Despite accumulating higher levels of ammonium, ryegrass exhibited slower physiological damage, suggesting a greater tolerance to ammonium toxicity, as also argued by Takano et al. (2019). This response could explain the delayed onset of senescence observed in this species.

## 4. Conclusions

The results of this study demonstrated a clear differential response between ryegrass (*Lolium multiflorum*) and barnyardgrass (*Echinochloa crus-galli*) to glufosinate ammonium, consistently observed across visual injury, chlorophyll fluorescence, and ammonium accumulation. Barnyardgrass exhibited a faster onset of symptoms, greater reductions in PSII quantum yield (Φ*_PSII_*), and more pronounced alterations in energy partitioning, indicating a higher sensitivity to the herbicide compared to ryegrass.

Although glufosinate ammonium is classified as a non-selective herbicide, our results confirm that its effectiveness can vary significantly among species, as previously reported (Takano et al., 2019). In this study, ryegrass showed a slower progression of injury and limited alterations in photosystem II functionality, suggesting a greater tolerance to glufosinate. These findings are consistent with previous reports describing reduced sensitivity of Lolium spp. to glufosinate, which has been associated with differences in herbicide absorption, translocation, and metabolism (Kumaratilake et al., 2002; Takano et al., 2019).

One of the most relevant findings of this study is that the differential sensitivity observed between species does not fully align with classical expectations based on photosynthetic metabolism. Previous studies have suggested that C4 species may exhibit lower sensitivity to glufosinate due to reduced photorespiration and lower ammonium accumulation (Wendler et al., 1990). However, in our study, the C4 species (barnyardgrass) showed a higher sensitivity than the C3 species (ryegrass), particularly in terms of early damage to PSII and faster senescence. This result suggests that factors beyond carbon metabolism, such as herbicide uptake, translocation efficiency, and detoxification capacity, may play a more decisive role in determining sensitivity.

Chlorophyll fluorescence proved to be a highly sensitive tool for detecting early physiological responses to glufosinate ammonium. Significant alterations in fluorescence parameters were observed within hours after application, prior to the appearance of visible symptoms. In barnyardgrass, the decrease in Φ*_PSII_* and the increase in basal energy dissipation (Φ*_NO_*) indicate structural damage to PSII and impaired electron transport. In contrast, ryegrass maintained relatively stable fluorescence parameters during the same period, supporting the hypothesis of delayed physiological damage in this species. These findings highlight the usefulness of fluorescence techniques for early detection of herbicide effects and for understanding their mode of action.

Ammonium accumulation further supports the observed differences between species. Although both species showed increased ammonium levels after herbicide application, barnyardgrass accumulated lower amounts than ryegrass, which is consistent with previous reports for C4 species (Wendler et al., 1990). However, despite this lower accumulation, barnyardgrass exhibited greater physiological damage, reinforcing the idea that ammonium accumulation alone does not fully explain herbicide sensitivity.

Overall, the results indicate that the response to glufosinate ammonium is species-dependent and cannot be explained solely by differences in photosynthetic metabolism. Instead, a combination of physiological and biochemical factors likely determines the final response. Understanding these mechanisms is essential for improving herbicide use efficiency and for designing management strategies aimed at delaying the evolution of resistance.

## Author’s contributions

Conceptualization of the manuscript and development of the methodology: O.S., M.S., S.S. and G.Q.;data collection and curation: O.S, G.Q, and S.S; data analysis: O.S., G.Q, and J.V.; data interpretation: G.Q. and J.V.; funding acquisition and resources: G.Q., and J.V.; project administration: G.Q., and J.V.; supervision: O.S. and G.Q; writing the original draft of the manuscript: M.S., S.S., and J.V.; writing, review and editing. All authors read and agreed to the published version of the manuscript.

## Data Availability

Data will be made available on request.

