## Supplemental data for "DIFFERENTIAL PHOTOSYNTHETIC RESPONSES TO GLUFOSINATE AMMONIUM IN TWO GRASS WEEDS: *Lolium multiflorum* AND *Echinochloa crus-galli*"

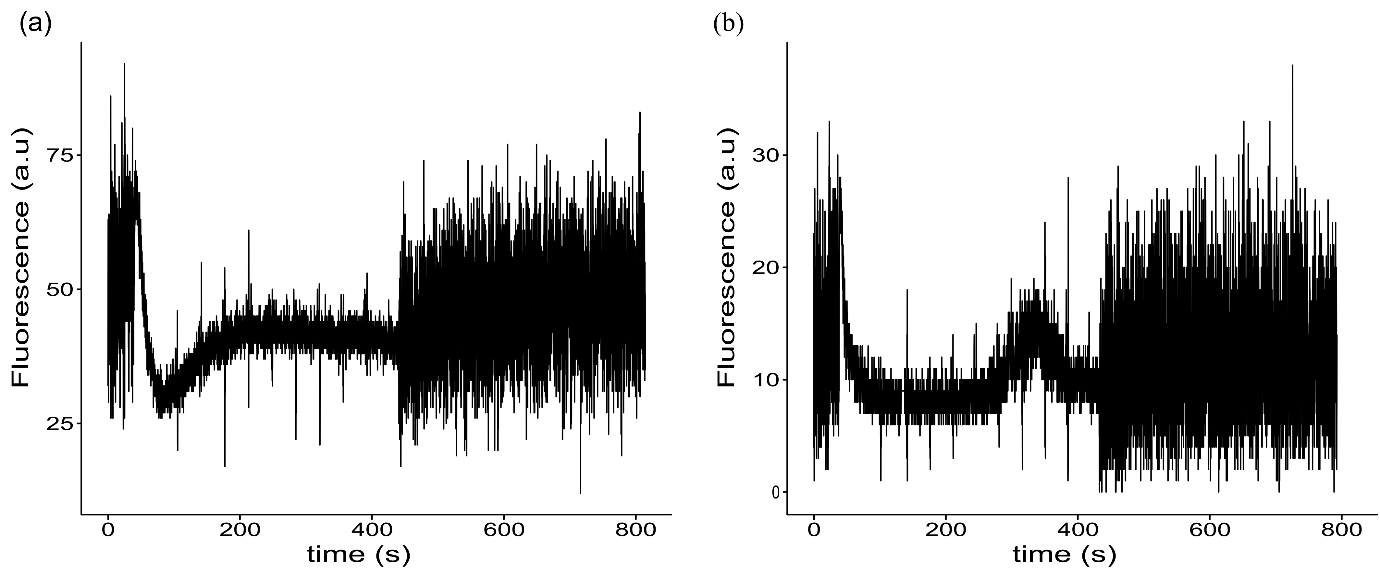

**Figure S1.** Chlorophyll fluorescence traces of photosystem II at 48 h after herbicide application.(a) *Lolium multiflorum* (L.)*,* (b) *Echinochloa crus-galli* (L.) Beauv.

Table S1. Análisis de varianza de
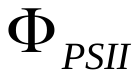
,
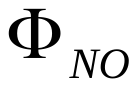
 y
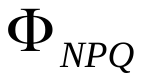

| Parameter | Effect | DFn | DFd | F | p | P<0.05 |
| --- | --- | --- | --- | --- | --- | --- |
| 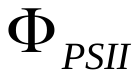 | Genotype | 2 | 54 | 153.4 | 5.3E-23 | * |
|  | Herbicide | 3 | 54 | 24.9 | 2.8E-10 | * |
|  | Time | 2 | 54 | 1.3 | 2.6E-01 |  |
|  | Genotype:Herbicide | 3 | 54 | 4.8 | 5.0E-03 | * |
|  | Genotype:Time | 2 | 54 | 0.6 | 5.1E-01 |  |
|  | Herbicide:Time | 6 | 54 | 1.6 | 1.4E-01 |  |
| 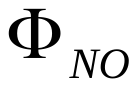 | Genotype | 2 | 54 | 55.9 | 6.9E-14 | * |
|  | Herbicide | 3 | 54 | 13.8 | 8.2E-07 | * |
|  | Time | 2 | 54 | 5.6 | 6.0E-03 | * |
|  | Genotype:Herbicide | 3 | 54 | 8.5 | 9.3E-05 | * |
|  | Genotype:Time | 2 | 54 | 5.4 | 7.0E-03 | * |
|  | Herbicide:Time | 6 | 54 | 4.3 | 1.0E-03 | * |
| 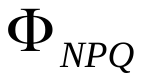 | Genotype | 2 | 54 | 48.6 | 8.3E-13 | * |
|  | Herbicide | 3 | 54 | 1.0 | 3.6E-01 |  |
|  | Time | 2 | 54 | 1.4 | 2.4E-01 |  |
|  | Genotype:Herbicide | 3 | 54 | 1.0 | 3.6E-01 |  |
|  | Genotype:Time | 2 | 54 | 1.8 | 1.6E-01 |  |
|  | Herbicide:Time | 6 | 54 | 0.9 | 4.5E-01 |  |
